# ASSET: A Framework for Decoding Aptamer Specificity of an Enriched Library by Next-Generation Sequencing of Experimental Samples

**DOI:** 10.1101/2025.08.27.672406

**Authors:** Maya Barros, Gayathri Kasirajan, Amara Jones, Aidan Schlichting, Dario Ruiz-Ciancio, Li-Hsien Lin, Chandan Narayan, Suresh Veeramani, Kristina W. Thiel, George Clare Kennedy, Isabel Darcy, William Thiel

## Abstract

The SELEX process to identify RNA and DNA aptamers relies on sequencing selection rounds to detect highly specific aptamers through patterns of aptamer accumulation or enrichment. However, this approach infers rather than quantify aptamer specificity. Here we present a novel strategy for directly quantifying aptamer specificity within enriched libraries termed **Aptamer Specificity Sequencing for Efficient Targeting (ASSET)**. The ASSET framework takes experimental samples and replicates testing the specificity of an aptamer library and prepares them for next-generation sequencing (NGS) with a *known internal reference sequence*. This enables robust data normalization, calculation of aptamer *specificity scores* with statistical significance, and the creation of *specificity profiles* of individual aptamers across multiple targets and non-targets. By integrating ASSET specificity scores with conventional selection round sequencing data, aptamers can be easily classified as true or false positives and negatives, allowing for easy separation of true positive aptamers. Compared to conventional methods for identifying aptamer candidates, such as measuring abundance or enrichment, ASSET specificity scores show a strong correlation with experimentally measured specificity. This supports ASSET as a more effective metric for selecting lead candidates following SELEX. ASSET is an easily implemented framework that accelerates the identification of highly specific aptamers, thereby expediting aptamer discovery for therapeutic and diagnostic applications.

## INTRODUCTION

Aptamers are short, single-stranded DNA or RNA molecules that bind target molecules with high specificity and affinity, making them a powerful technology for a wide range of biomedical applications, including diagnostics^1^, imaging^2^, therapeutics^3^, and delivery^4^. Their clinical relevance continues to grow, with several aptamers currently undergoing clinical trials for both diagnostic and therapeutic purposes.^5^ Notably, two aptamer-based therapeutics have received FDA approval, the most recent in 2023 for the treatment of an ocular disorder.^7^

Aptamer discovery typically uses Systematic Evolution of Ligands by EXponential enrichment (SELEX), an iterative process that begins with libraries of aptamers and enriches for sequences with high affinity for a specific target.^6, 7^ Since its introduction in 1990, SELEX has undergone numerous refinements and innovations to improve the specificity and efficiency of paring down complex aptamer libraries.^8-10^ With the implementation of next-generation sequencing (NGS) with aptamer selections^11^, researchers can now analyze millions of aptamer sequences across iterative selection rounds, enabling computational approaches to identify patterns of enrichment and infer candidate aptamers or motifs with potential target specificity.^12, 13^ Recent algorithms such as RaptGen^14^ and AptaDiff^15^ leverage these data with machine learning to enhance generative aptamer discovery.

However, it is well established that the most abundant or enriched sequences are not always the most specific or highest-affinity aptamers.^16-18^ Factors such as suboptimal selection conditions or amplification bias introduced during PCR in the SELEX process can result in the accumulation of low affinity sequences.^10^ Consequently, sequencing data from selection rounds provides limited insight into which aptamers exhibit the greatest specificity for a target relative to non-targets. This limitation becomes increasingly complex when multiple preferred targets and non-targets are involved.

To address these limitations, we present **ASSET (Aptamer Specificity Sequencing for Efficient Targeting)**, a robust, and easy-to-implement NGS-based strategy for identifying high-specificity aptamers. Unlike conventional SELEX sequencing, which only infers specificity from abundance and enrichment patterns, ASSET incorporates an internal reference sequence to normalize data across replicates and conditions, allowing the calculation of specificity scores for every aptamer in a library. This approach allows for the identification of truly specific candidates from false positives. This enables informed candidate selection without the need to sequence the SELEX selection rounds.

## RESULTS

### ASSET - Aptamer Specificity Sequencing for Efficient Targeting

To address the challenge of comparing NGS results across experimental samples and replicates in aptamer specificity assays, we developed the ASSET framework (**Figure 1**). Unlike conventional sequencing of SELEX selection rounds, which infers specificity from patterns of aptamer accumulation and enrichment, the ASSET framework introduces a fundamentally different strategy. Experimental samples evaluating an aptamer library’s specificity for targets and non-targets are prepared with a *known internal reference sequence*. The internal reference sequence enables precise normalization of NGS read counts across conditions and direct calculation of a *specificity score*. Specificity scores are the fold specificity of an aptamer for a target over non-target. Incorporation of replicates for target and non-target samples allows for statistically significance to be assessed. A compilation of specificity scores allows creation of a robust *specificity profile* of aptamers within the enriched library across any number of targets and non-targets.

**Figure 1.**
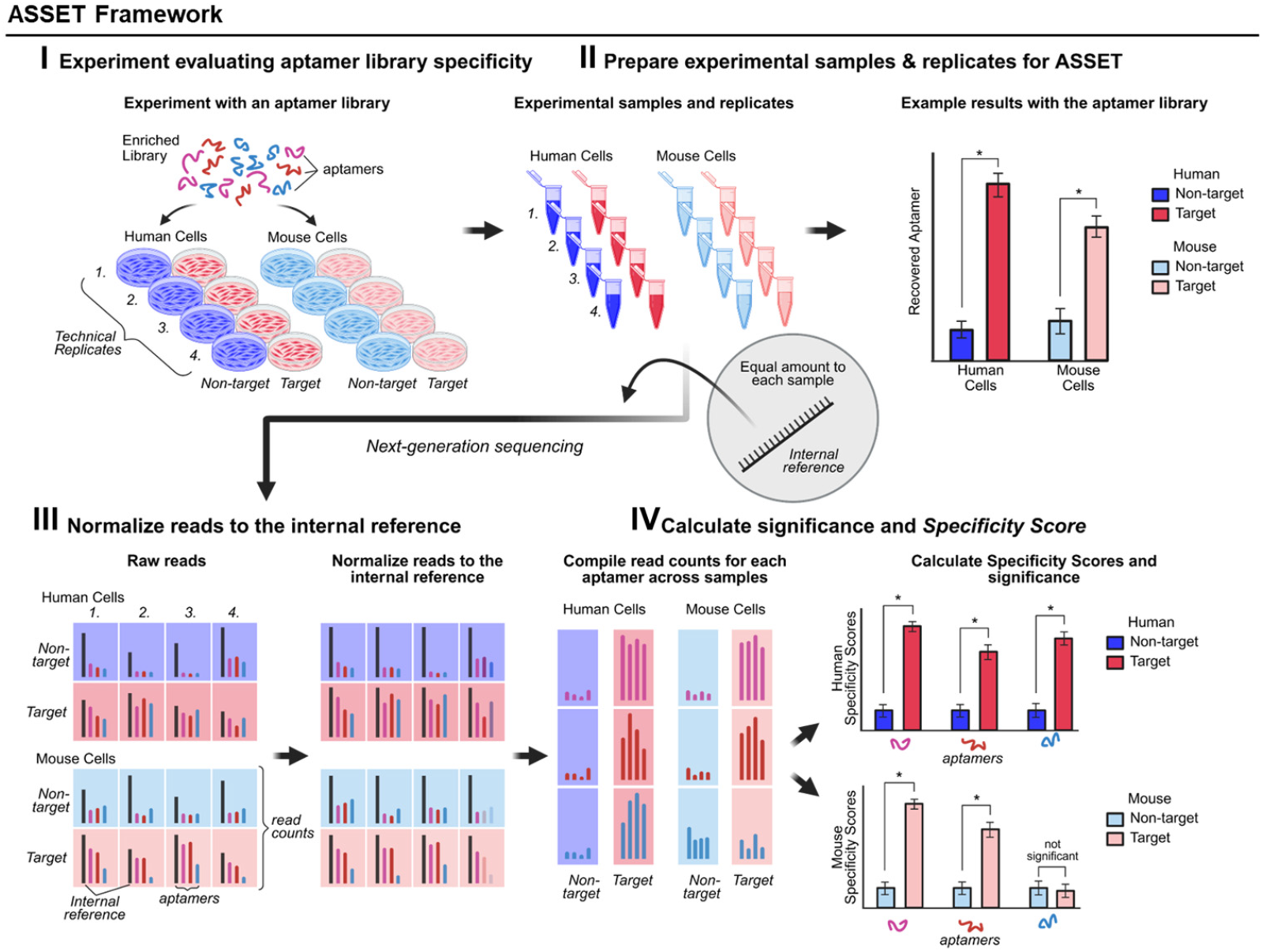
The ASSET Framework. (I) Samples and replicates from experiments evaluating an enriched aptamer library’s specificity are (II) prepared for NGS with an equal amount of an *internal reference*. (III) Read counts across samples and replicates are normalized to the internal reference. (IV) Read counts for each aptamer (3 shown) are compile for each of the four technical replication (4 bars per aptamers). From the normalized read counts, a *specificity score* for each aptamer can be calculated and the significance of specificity determined. These data provide a specificity profile for each aptamer detected within the enriched library across any number of targets and non-targets.

### SELEX enrichment of internalizing aptamers specific for primary human and mouse VSMCs

We applied ASSET to an aptamer library enriched for internalizing aptamers that target human coronary artery vascular smooth muscle cells (VSMCs), aiming to improve specificity relative to coronary artery endothelial cells. This selection built upon aptamers previously identified using immortalized rodent vascular cell lines^19^, which demonstrated therapeutic promise in mouse and pig models of vascular disease^20-23^. However, these aptamers have capacity for internalization into primary human VSMCs (**Supplemental Figure 1**). Our goal herein was to identify aptamers with a significantly higher capability for internalizing into human VSMCs over endothelial cells that also cross-react well with mouse VSMCs. These aptamers would support future targeted therapeutics for human vascular diseases driven by VSMC pathologies while allowing *in vivo* evaluation with mouse models of vascular disease.^24, 25^

To enhance the translational relevance of these aptamers, we introduced several strategic modifications to the Cell-Internalization SELEX protocol (**Figures 2a and 2b**). First, we employed optimized *in vitro* transcription to generate a 2’-O-methyl (OMe) N25 RNA aptamer library^26^, a chemical modification known to reduce immunogenicity compared to 2’-fluoro chemistry.^27, 28^ To enrich for aptamers with human specificity while preserving compatibility for *in vivo* evaluation in mouse models, we adapted the Toggle SELEX method^29^ to a cell-based format by alternating selection between human and mouse vascular cells. To ensure robust selection of internalizing aptamers, we replaced the conventional high-salt wash with an RNase cocktail that effectively degrades surface-bound but not internalized OMe-modified aptamers (**Supplemental Figure 2a**). In the final selection rounds, we shortened the incubation time to 15 minutes to favor aptamers that target high-turnover cell-surface proteins. Finally, we conducted the initial rounds in triplicate to reduce the risk of losing potential aptamer candidates during early stage of SELEX. These refinements were applied over nine selection rounds and are summarized in **Supplemental Figure 2b**.

**Figure 2.**
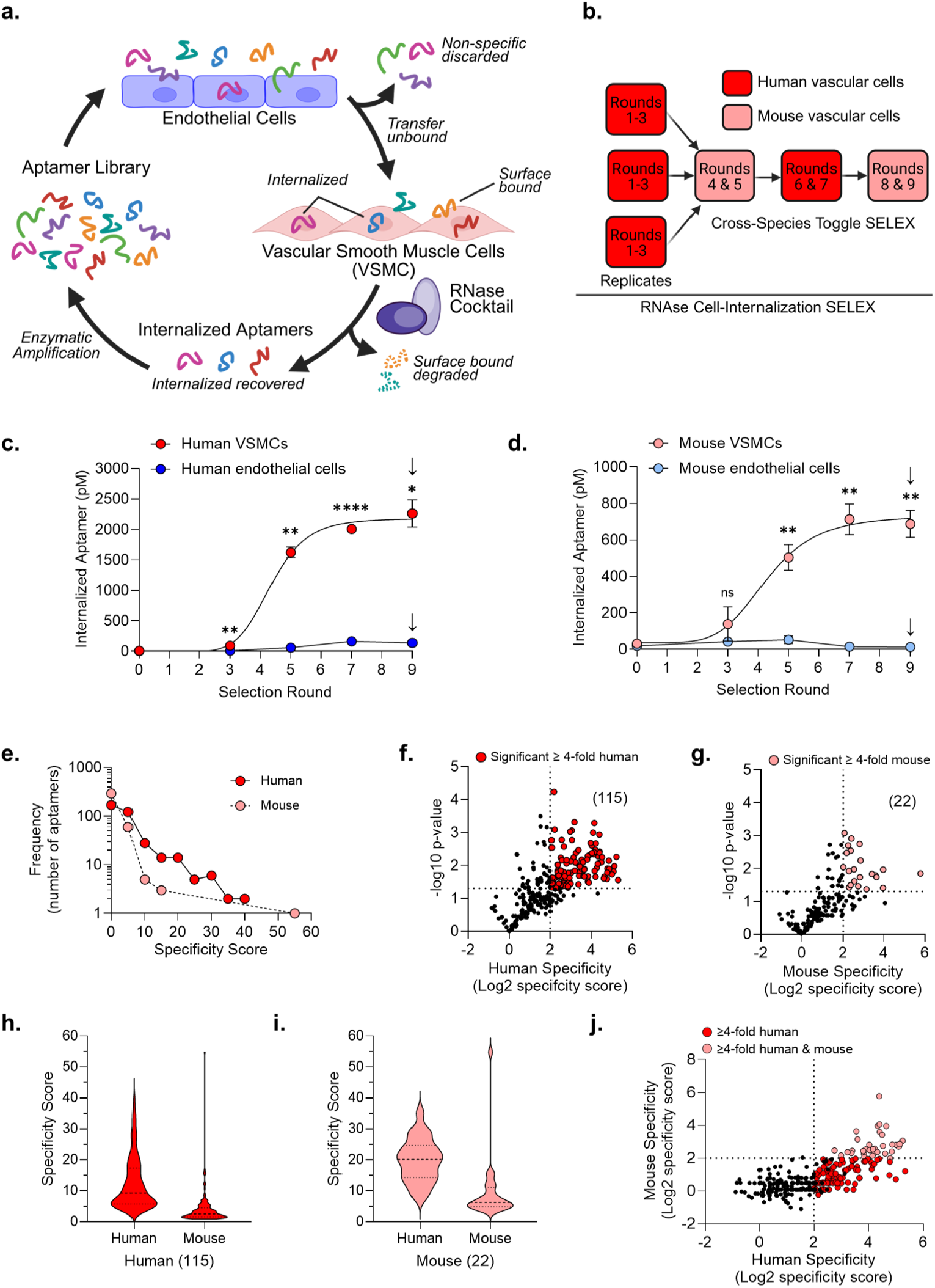
Application of the ASSET framework to a Cell-Internalization SELEX for internalizing aptamers specific for both human and mouse primary VSMCs. (**a**) A schematic of the Cell-Internalization SELEX selection round strategy to generate VSMC internalizing aptamers. (**b**) The overall Toggle SELEX strategy with replicates applied to the early selection rounds to generate aptamers specific for both human and mouse VSMCs. Quantitative PCR data from an internalization experiment evaluating the specificity of the VSMC aptamer library selection rounds for (**c**) human vascular cells, and (**d**) mouse vascular cells. N = 3-4, 2-way ANOVA with multiple comparisons, ^*^ p < 0.05, ^**^ p < 0.005, ^****^ p < 0.0001, ns = not significant. ↓ = samples and replicates prepared for ASSET. (**e**) Distribution (Frequency, number of aptamers) of the human and mouse ASSET specificity scores for VSMCs. Volcano plots of (**f**) human and (**g**) mouse ASSET specificity scores for VSMCs versus specificity score significance determined by an unpaired Student’s T-test (-log10 p-value). 115 aptamers exhibited significant specificity for human VSMCs greater than four-fold. 22 aptamers exhibited significant specificity for mouse VSMCs greater than four-fold. Violin plots of the (**h**) 115 and (**i**) 22 aptamer’s specificity scores for human and mouse VSMCs. (**j**) Relationship between aptamer human and mouse specificity scores (Log2 Specificity Score).

We evaluated the starting aptamer library and selection rounds for specificity and internalization capacity against both human and mouse vascular cells. By round nine, the library demonstrated robust internalization into VSMCs over endothelial cells in both species, with a notably stronger and earlier signal detected with human VSMCs (**Figure 2c,d**). Specificity for human VSMCs was evident by round three and continued to increase through round nine, indicating consistent enrichment of aptamers with human VSMC preference (**Figure 2c**). In contrast, mouse VSMC specificity emerged more gradually, with the most pronounced shift occurring between rounds three and five, followed by only modest gains thereafter (**Figure 2d**). These interspecies differences in library behavior are likely attributable to the Toggle SELEX protocol, which was initiated with human vascular cells, with mouse cells introduced beginning in round four.

### ASSET evaluation of VSMC internalizing aptamer cross-species specificity

To quantify specificity with high precision, we applied the ASSET framework to the technical replicates from the round nine internalization assay in Figure 2c and 2d, enabling direct comparison of aptamer specificity for human and mouse VSMCs. Rather than sequencing rounds, we performed NGS on aptamers recovered from target and non-target cells as depicted in Figure 1. Across all experimental replicates (i.e., target and non-target, murine and human cells), ASSET data revealed between 262 and 597 unique aptamer sequences, representing 255,071 reads (**Supplemental Figure 3a**). We examined the distribution of these aptamer sequences across replicates (**Supplemental Figure 3b**) and identified 363 unique aptamers that were consistently present in at least three of the four replicates of human VSMC. Focusing on this subset, we assessed the distribution of specificity scores for human and mouse VSMCs. The specificity scores indicate that the Toggle SELEX process successfully enriched for aptamers with specificity for both human and mouse VSMCs, with a clear predominance toward human (**Figure 2e**).

To identify the top aptamer candidates, we selected those with specificity scores exceeding a four-fold threshold and that also showed statistical significance by a Student’s t-test. This threshold was selected to exceed the specificity observed in first-generation 2’-fluoro-modified VSMC aptamers (**Supplemental Figure 1**), ensuring that only highly specific sequences were considered. Volcano plots were applied to visualize log2-transformed specificity scores against statistical significance (-log_10_ p-values), thereby highlighting aptamers that meet criteria for specificity and significance. From the human ASSET data, 115 aptamers demonstrated significant four-fold specificity for human VSMCs (**Figure 2f**), while 22 aptamers met the same threshold for mouse VSMCs (**Figure 2g**).

The 115 aptamers that exhibit significant and greater than four-fold human-specificity exhibited a mean specificity score of 12.41 (±0.79 SEM) and a median of 9.3, while mouse specificity averaged 4.02 (±0.53 SEM) with a median of 2.51 **(Figure 2h, Supplemental Figure 3c**). The 22 mouse-specific aptamers showed even higher human specificity, with a mean score of 19.85 (±1.50 SEM) and a median of 20.13, alongside a mean mouse specificity score of 9.66 (±2.28 SEM) and a median of 6.25 (**Figure 2i, Supplemental Figure 3d**).

We further examined cross-species reactivity by generating a *specificity profile* by plotting human and mouse VSMC specificity scores against each other (**Figure 2j**). This analysis revealed a strong and statistically significant Pearson’s correlation (r = 0.72, p <0.0001), indicating that a subset of aptamers specific to one species also cross-react with the other. This statistically significant relationship between human and mouse specificity scores was shown to also be strong Spearman’s correlation (r = 0.63, p <0.0001), but moderate by Kendall’s correlation (r = 0.47, p <0.0001). Notably, the ASSET specificity profile effectively identifies a subset of cross-reactive aptamers, highlighting its utility in guiding aptamer selection across multiple targets and non-targets.

### Comparison between ASSET and NGS of the SELEX selection rounds

To evaluate the depth and effectiveness of the ASSET framework, we compared our ASSET data to NGS data acquired from directly sequencing the selection rounds. Following standard naming conventions, the individual VSMC aptamers are labeled based on their rank order in round nine, determined by read count.

Conventional SELEX-NGS was performed by sequencing the starting aptamer libraries and all nine selection rounds. These data identified 84,251,647 unique aptamer sequences comprising 157,360,021 total reads (**Supplemental Figure 4a**). Sequence enrichment analysis revealed that 50% of library enrichment occurred between rounds three and four, with maximal enrichment achieved by round six (**Figure 3a**).

**Figure 3:**
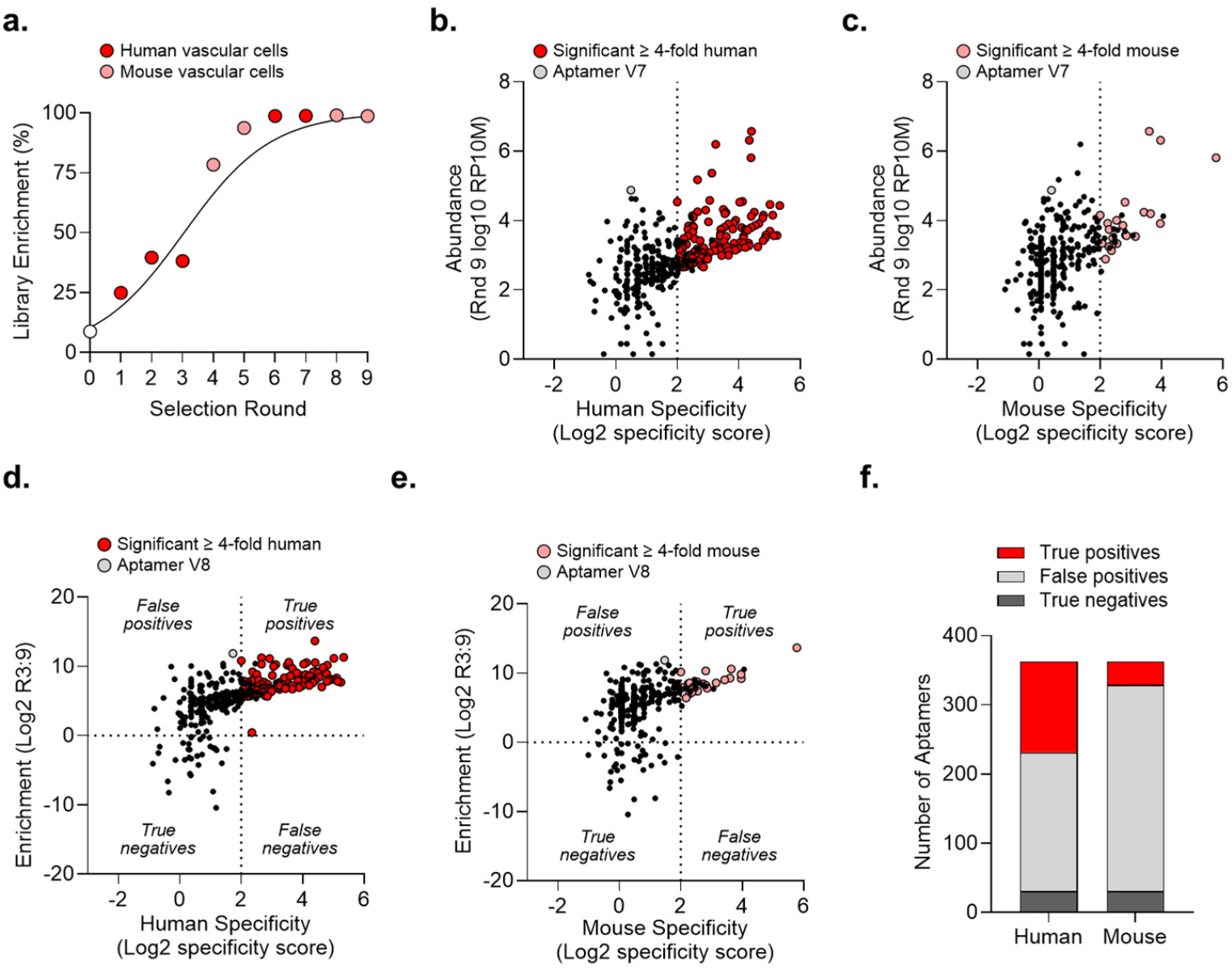
Comparison and integration of ASSET specificity scores with SELEX selection round enrichment data. (**a**) Enrichment of the aptamer library as determined by the proportion of NGS unique reads to total reads for each selection round. % Library Enrichment = 100^*^(1 - (Unique reads/Total reads)). Relationship of ASSET (**b**) human and (**c**) mouse specificity scores (log2 specificity score) between the log10 reads per 10 million aptamer abundance within round nine (Rnd 9 log10 RP10M) as determine by Pearson’s correlation. Relationship of ASSET (**d**) human and (**e**) mouse specificity scores (log2 specificity score) between the aptamer enrichment from round three to round nine (log2 R3:9). Aptamers with positive log2 enrichment and greater than four-fold specificity are classified as true positives. Aptamer with positive log2 enrichment and less than four-fold specificity are classified as false positives. Aptamer with negative enrichment and less than four-fold specificity are classified as true negatives. Aptamers with negative enrichment and greater than four-fold specificity are classified as false negatives. (**f**) Distribution of aptamers determined to be true positives, false positive, true negatives, and false negatives for human and mouse VSMCs based on specificity scores and round three to nine enrichment.

To evaluate the depth of ASSET data coverage, we identified the lowest ranked aptamer by abundance from round nine sequencing data that was also detected by ASSET. The ASSET dataset included aptamers ranked as low as 2,560th in round nine. Among aptamers with significant four-fold specificity, the 115 human specific aptamers included sequences ranked as low as 332nd, while the 22 mouse specific aptamers included one ranked 238th. These findings indicate that ASSET provides specificity scores for approximately the top 2,500 most abundant aptamers within round nine. Furthermore, the most specific aptamers tend to be found within the top 200 to 300 by abundance.

Next, we compared ASSET specificity scores to aptamer abundance and enrichment metrics, which are commonly used to choose candidate sequences (**Figures 3b and 3c**). When evaluating ASSET specificity scores against aptamer abundance, we observed a significant but weak linear Pearson’s correlation for both human (r = 0.18, p = 0.0007) and mouse (r = 0.28, p <0.0001) VSMCs. However, the rank order correlation as determined by Spearman’s correlation (human r = 0.61, p<0.0001; mouse r = 0.44, p <0.0001) and Kendall’s correlation (human r = 0.45, p < 0.0001; mouse r = 0.44, p<0.0001) better (**Supplemental Figure 4b**). While some of the most abundant aptamers exhibited high specificity, others did not. For example, among the top ten most abundant aptamers in round nine (**Supplemental Figure 4c**), three showed greater than twenty-fold human and ten-fold mouse specificity, while four had specificity scores less than four-fold for both species. Notably, aptamer V7, ranked 7th in abundance, exhibited poor specificity scores for both human (1.41) and mouse (1.33) VSMCs (annotated in grey in **Figure 3b, c**). These findings suggest that while abundance may help identify some high-specificity candidates, it is not a reliable predictor of aptamer specificity.

We also evaluated changes in aptamer enrichment to assess cross-species specificity, focusing on differences between round three, before mouse vascular cell introduction, and round nine, the final round exposed to mouse cells. Comparing ASSET specificity scores to round three to nine enrichment values (**Figures 3d and 3e**), we observed significant and stronger Pearson’s correlation (**Supplemental Figure 4b**; human r = 0.60, p <0.0001; mouse r = 0.40, p <0.0001), Spearman’s (human r = 0.70, p <0.0001; mouse r = 0.47, p <0.0001) and Kendall’s correlation (human r = 0.53, p <0.0001; mouse r = 0.34, p<0.0001). These data suggest aptamer enrichment appears to serve as a more reliable metric to identify specific aptamers than abundance. However as with abundance, the top ten most enriched aptamers included both highly specific and poorly specific aptamers (**Supplemental Figure 4c**). For instance, aptamer V8, the second most enriched sequence and eighth most abundant (annotated in grey in **Figure 3c, d**), showed low specificity scores of 3.31 for human and 2.78 for mouse VSMCs. These results indicate that enrichment, like abundance, while still not a reliable metric for predicting aptamer specificity may be better than aptamer abundance.

### Integration of ASSET specificity scores and selection round sequencing data enables aptamer classification

By integrating ASSET specificity scores with selection round enrichment data, we classified aptamers into four categories: true positives, true negatives, false positives, and false negatives. *True positives* were defined as aptamers that were positively enriched and exhibited specificity for VSMCs (target) of at least four-fold over endothelial cells (non-target). *True negatives* were negatively enriched and showed specificity of four-fold or less. *False positives* were positively enriched but lacked sufficient specificity, while *false negatives* were negatively enriched yet demonstrated strong specificity. This classification provides insight into the efficiency of the SELEX process, where a high proportion of true positives is indicative of successful selection.

For human-specific aptamers, we observed a large proportion of false positives, followed by a nearly equal number of true positives and a smaller fraction of true negatives **(Figure 3e**). In contrast, mouse-specific aptamers showed a higher proportion of false positives and a lower number of true positives relative to human-specific aptamers, while the fraction of true negatives was similar to human data (**Figure 3f**). Notably, no false negatives were identified in either dataset.

These results suggest that the number of ideal aptamers, true positives with cross-species reactivity, is relatively small compared to the total pool of sequenced aptamers in a converged library. The prevalence of false positives highlights the limitations of relying solely on abundance or enrichment metrics for aptamer selection. Importantly, these findings underscore the value of incorporating quantitative specificity data, such as that provided by ASSET, to improve the accuracy and reliability of post-SELEX candidate prioritization.

### Identification and empirical evaluation of top ASSET candidates

To identify lead VSMC aptamers with strong cross-species reactivity, we determine a threshold based on specificity scores two standard deviations above the mean detected across all 363 aptamers (**Supplemental Figure 5a and 5b**). The 363 aptamers identified by ASSET exhibited a mean specificity score of 5.49 (±6.84 SD), with mouse specificity averaged 2.15 (±3.46 SD). This resulted in mean plus two standard deviation thresholds of 19.18 for human data and 9.07 for mouse data. Based on these criteria, we prioritized aptamers demonstrating greater than 20-fold specificity for human and 10-fold specificity for mouse. Due to the limited number of aptamers meeting these criteria, we also included aptamer V28, which, despite not reaching statistical significance for mouse specificity, demonstrated over ten-fold mouse and twenty-fold human specificity. Additionally, due to the overall lower specificity observed for mouse VSMCs, we ranked candidates based on mouse specificity. This approach yielded six lead aptamers (**Figures 4a and 4b**).

**Figure 4.**
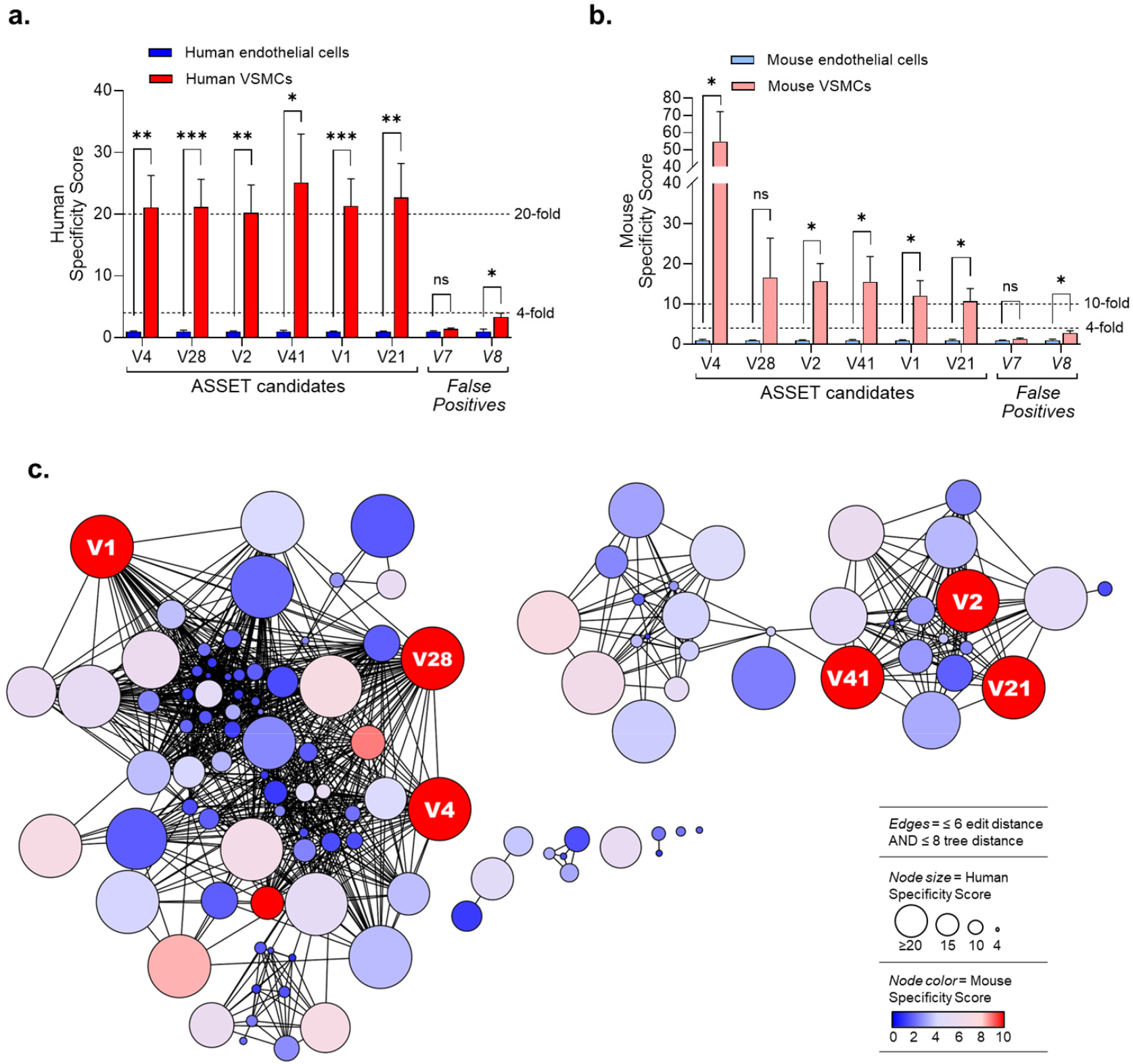
Identification of top cross-species reactive VSMC aptamers using ASSET specificity scores. VSMC aptamers with (**a**) greater than twenty-fold specificity for human VSMCs, and (**b**) greater than ten-fold specificity for mouse VSMCs. False positive control aptamers, V7 and V8, exhibit less than four-fold specificity scores from human and mouse VSMCs. (**c**) AptamerRunner clustering of 115 VSMC aptamers by edit distance six and tree distance eight. Nodes denote individual aptamers with node size determine by human VSMC specificity scores and node color determined by mouse VSMC specificity scores. Nodes representing the top candidate aptamers are labeled.

To assess sequence and structural diversity among the candidates, we clustered the 115 aptamers exhibiting greater than four-fold human VSMC specificity using an edit distance threshold of six and a tree distance threshold of eight and noted the location of the top candidates (**Figure 4c**). AptamerRunner revealed two major clusters that contained the majority of the top candidates. From these data, we selected three candidates within the first major cluster (V4, V28, and V1), and two candidates from the second major cluster (V2, V41). We also included V7 and V8 as negative controls, representing false positives identified by abundance and enrichment but lacking strong ASSET specificity score.

The lead candidates and negative controls (**Figure 5a**) were tested for internalization into human and mouse vascular cells. The top ASSET-identified aptamers demonstrated excellent specificity for VSMCs over endothelial cells in both species (**Figures 5b and 5c**), with human specificity ranging from 12.7-to 44-fold (**Figure 5d**) and mouse specificity from 21.7-to 72-fold (**Figure 5e**). Notably, the observed mouse specificity aligned more closely with the ASSET-based ranking than with the candidates’ rank order from round nine of the SELEX process. The negative control aptamers V7 and V8 showed no significant specificity for human VSMCs over endothelial cells and only modest mouse specificity (2.6-to 3.8-fold), confirming their classification as false positives. These results validate ASSET specificity scores as reliable predictors of aptamer specificity.

**Figure 5:**
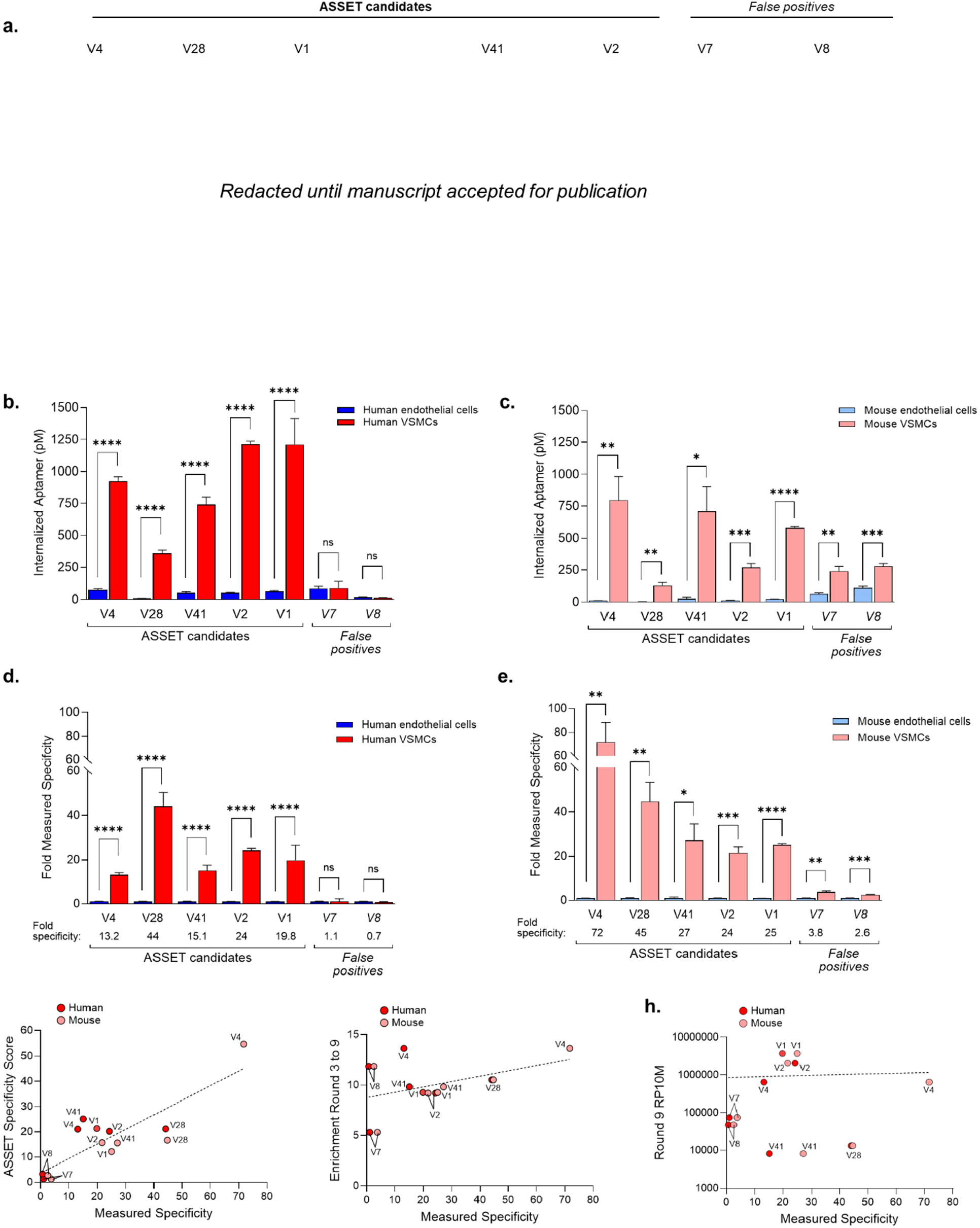
Evaluation of top cross-species reactive VSMC aptamers. (**a**) Predicted secondary structure of VSMC aptamer (V4, V28, V1, V41, V2), and false positive control aptamers (V7, V8). Aptamers grouped together are from the same cluster of aptamers related by sequence and structure. Quantitative PCR of internalized aptamer from (**b**) human and (**c**) mouse vascular cells treated with candidate and control aptamers. Fold specificity of candidate and control aptamers for (**d**) human and (**e**) mouse VSMCs over endothelial cells. N=4, Multiple unpaired T-tests. ^*^ p < 0.05, ^**^ p < 0.01, ^***^ p < 0.001, ^****^ p <0.0001. Relationship between measure specificity and (**f**) ASSET specificity scores, (**g**) aptamer enrichment from round three to round nine, and (**h**) aptamer abundance in round nine (Round 9 RP10M).

To quantify the relationship between predicted and observed specificity, we plotted ASSET specificity scores against experimentally measured specificity (Figure 5f) and assessed their correlation (**Supplemental Figure 6b**). Pearson’s correlation revealed a strong linear relationship (r = 0.84, *p* = 0.001), indicating that ASSET scores closely reflect real-world specificity. In contrast, Spearman’s correlation showed a moderate but significant association (ρ = 0.57, *p* = 0.35), while Kendall’s correlation indicated a weak and statistically insignificant relationship. For comparison, we also evaluated measured specificity against aptamer enrichment from rounds three to nine (**Figure 5g**) and abundance in round nine (**Figure 5h**). Both metrics, regardless of the correlation method used, exhibited weak to very weak and non-significant associations with measured specificity.

Collectively, these findings demonstrate that ASSET is a superior method for identifying high-specificity aptamers compared to conventional ranking by abundance or enrichment, summarized in **Supplemental Figure 6**. ASSET enables accurate prediction of aptamer performance, streamlining candidate selection and improving the efficiency of post-SELEX validation.

## DISCUSSION

Our primary goal with this study was to devise a novel NGS-based approach to enhance the ability to identify specific aptamers without laborious post-SELEX screening of candidates. ASSET represents a major advancement over traditional analysis of selection rounds that infer aptamer specificity based on patterns of aptamer accumulation during SELEX. In contrast, ASSET directly quantifies aptamer specificity within enriched libraries using NGS, based on experiments explicitly designed to evaluate specificity. By incorporating experimental replicates, ASSET also enables statistical testing of specificity across multiple target and non-target conditions. Our results show that ASSET alone, rather than sequencing data from selection rounds, is not only sufficient, but also ideal for identifying aptamers with high specificity.

Using ASSET, we identified five aptamers with desired specificity profiles for both human and mouse VSMCs. We further validated ASSET’s accuracy by showing that two other aptamers, despite being highly abundant or enriched in the final selection, exhibited limited specificity for human and mouse VSMCs, consistent with their low ASSET specificity scores. These findings highlight ASSET’s ability to reveal comprehensive specificity profiles across multiple targets and non-targets. Because ASSET requires only the inclusion of a reference sequence during sequencing amplicon preparation, it can be seamlessly integrated into existing aptamer selection workflows, including retrospective analysis of past experiments when samples are available.

We identified only one study that attempts to quantify aptamer interactions between targets and non-targets using NGS data.^30^ The study by Pleiko *et al*. applied the RNA-seq analysis tools edgeR^31^ to assess differential aptamer binding to target and non-target cells from a Cell-SELEX. While the concept is compelling, we believe the authors overlooked a critical aspect of edgeR related to data normalization. From edgeR guidelines, “normalization issues arise only to the extent that technical factors have sample-specific effects,” with examples including “varying sequencing depths.” When comparing the amount of total aptamer bound to target and non-target samples, the factor of normalization due to sample-specific effects, including varying NGS sequencing depth, are of critical importance. This problem is less prominent for RNA-seq when recovering RNA from cells, which includes common genes that constitute a biological internal standard. Therefore, addressing challenges related to normalization is essential for accurately normalizing data attained from experiments testing an aptamer library’s specificity. The ASSET framework solves this problem by the addition of a known internal reference sequence.

While ASSET data are promising, there remain future opportunities for refinement. For instance, our data show that the internal reference sequence accounted for approximately 89 percent of sequenced reads. Although this coverage was sufficient to assess the specificity of the top 2,500 ranked aptamers, the high proportion of reference reads suggests room for improvement in balancing reference and library sequences. Future work will explore how this ratio can be adjusted without compromising accuracy or reproducibility. Next, some differences were observed between ASSET specificity scores and the measured specificity of the VSMC aptamers. These differences likely stem from interactions within the enriched library, whereby aptamers may compete or cooperate in ways that influence binding/internalization behavior. Such interactions are not present when single aptamers are tested empirically. Nevertheless, the strong correlation between ASSET specificity scores and experimental results, especially the consistent ranking of mouse VSMC specificity, supports ASSET’s accuracy, reliability and utility.

Future applications of ASSET could address and long-standing limitation of SELEX: the challenge of identifying aptamers from *in vitro*-enriched libraries that retain functionality in more complex systems, such as *in vivo* animal models of disease and patient-derived tissues or organoids.^32^ By enabling direct testing of enriched libraries against these more complex models, ASSET offers a powerful strategy to identify aptamers with the greatest potential for clinical translation. This capability opens a new path for aptamer discovery, where candidates selected through reproducible *in vitro* methods can be rapidly evaluated in biologically relevant contexts. Ultimately, ASSET has the potential to accelerate the development of aptamers for therapeutic and diagnostic applications by bridging the gap between SELEX and real-world biological complexity.

## CONCLUSION

ASSET represents a paradigm shift in the identification of highly specific aptamers from enriched libraries, replacing the 35-year reliance on sequencing selection rounds with a direct and quantifiable approach that sequences experiments evaluating library specificity.

## METHODS

### 2’-OMe-pyrimidine modified RNA aptamer library

The Sel2 aptamer library with 25 nucleotide random region (Sel2N25) ssDNA template oligonucleotide was synthesized at a 1 µM scale (IDT, Coralville IA): 5′-[UC]GGGCGAGTCGTCTG N25 CCGCATCGTCCTCCC-3′, where [UC] denotes 2′-O-methyl (OMe) modification to improve *in vitro* RNA transcription product.^26^ The Sel2 5′ primer (5′-TAATACGACTCACTATAGGGAGGACGATGCGG-3′) and the Sel2 3′ primer (5′-[UC]GGGCGAGTCGTCTG-3′) were used for amplification, with the 3′ primer also containing the [UC] OMe modification. The aptamer template oligonucleotide was extended following previously described protocols. *In vitro* transcription was performed using a Y639F mutant T7 RNA polymerase^33^ in the presence of OMe-modified pyrimidine nucleotides, with the reaction supplemented with 1.5 mM MnCl_2_ to enhance transcription efficiency. The resulting RNA transcripts were purified as previously described.^26^

### Cells

Human coronary artery vascular smooth muscle cells (ATCC, PCS-100-021) were cultured using the corresponding media kit (ATCC, PCS-100-042) and vascular cell basal medium (ATCC, PCS-100-030). Human coronary artery endothelial cells (ATCC, PCS-100-020) were maintained under similar conditions using the appropriate endothelial cell media kit (ATCC, PCS-100-041) and vascular cell basal medium. Mouse aortic VSMCs derived from C57BL/6 mice (Cell Biologics, C57-6080) were cultured using the Cell Biologics VSMC media kit (M2268), while mouse aortic endothelial cells from the same strain (Cell Biologics, C57-6052) were maintained with the corresponding endothelial cell media kit (M1168). Human and mouse vascular cells were cultured at 37°C in a humidified incubator under 5% CO_2_, propagated per each company’s instructions. Cells were lot-matched and used between passage six to eight for SELEX selection rounds and all experiments.

### RNase Cell-Internalization SELEX

Experimental conditions applied during SELEX are summarized in **Supplemental Figure 2b**. Cells were plated in 10 cm culture dishes at a density of 30,000–35,000 cells/cm^2^ and maintained at 37°C in a humidified incubator with 5% CO_2_ for a minimum of 24 hours. Aptamer RNA was folded in Binding Buffer containing 150 mM NaCl, 2 mM CaCl_2_, and 20 mM HEPES by heating at 94°C for 5 minutes, followed by passive cooling on a heat block for 45 minutes and placement on ice for at least 5 minutes. To degrade extracellular RNA, a nuclease cocktail was prepared using 5,000 Units/mL RNase T1 (ThermoScientific, FEREN0541) and 2,000 gel units/mL Micrococcal Nuclease (NEB, M0247S) in cell culture media. After incubation with the RNase cocktail, internalized aptamer RNA was recovered by TRIzol extraction supplemented with 20 mg/mL glycogen. The purity of the extracted RNA was assessed using a Nanodrop spectrophotometer and cleaned if necessary. Recovered aptamer RNA was reverse transcribed using Superscript IV (Thermo Fisher Scientific, 18090050) and amplified by PCR using Q5 Hot Start High-Fidelity DNA Polymerase (NEB, M0493S) to generate dsDNA, which was quantified by Nanodrop and subsequently used for *in vitro* transcription to generate the aptamer library. Transcribed aptamer RNA was purified as previously described and quantified by Nanodrop.

### Internalization assay

Human and mouse vascular cells were seeded at a density of 100,000 cells per well in 12-well plates and cultured for a minimum of 24 hours prior to treatment. Aptamer RNA was folded and diluted to a final concentration of 150 nM for aptamer library or 50 nM for aptamer candidates and controls in culture media specific to each cell type, supplemented with 100 ng/mL yeast tRNA. To minimize nonspecific binding, cells were pre-blocked with yeast tRNA for 15 minutes, followed by incubation with aptamer for 30 minutes. After treatment, cells were washed with ice-cold culture media and exposed to the RNase cocktail for 5 minutes at 4°C to degrade surface bound and unbound aptamer. Cells were then lysed using TRIzol reagent containing 20 mg/mL glycogen, and aptamer RNA was recovered by organic extraction using phase lock heavy tubes. The quantity of recovered aptamer RNA was assessed by quantitative reverse transcription PCR (RT-qPCR) using a two-step protocol: cDNA synthesis was performed with SuperScript IV reverse transcriptase, followed by amplification with iQ SYBR Green Supermix (Bio-Rad). Reactions were run on a QuantStudio 3 Real-Time PCR System, using the Sel2N25 OMe round 0 aptamer library as a standard across both RT and qPCR steps. Water controls were included for both RT and qPCR to monitor for nonspecific amplification.

### ASSET NGS amplicon preparation

To generate the ASSET amplicon for NGS, RT-PCR was performed in 10 µL reaction volumes using Illumina-compatible reverse primers and barcoded forward primers. Each sample was spiked with 10 pmol of a chemically synthesized Control-2F aptamer (TriLink). The Illumina reverse primer sequence was 5′-CAAGCAGAAGACGGCATACGAGATTCGGGCGAGTCGTCTG-3′, and the barcoded forward primer included the Illumina adapter sequence followed by a custom eight nucleotide barcode and the aptamer-specific region: 5′-AATGATACGGCGACCACCGAGATCTACACTCTTTCCCTACACGACGCTCTTCCGATCT-BARCODE-GGGAGGACGATGCGG-3′. Reverse transcription was carried out using SuperScript IV (ThermoFisher Scientific), and amplification was performed using Q5 Hot Start High-Fidelity DNA Polymerase (NEB) for 15–18 cycles. PCR products were evaluated for the correct amplicon size on 4% E-Gel EX agarose gels (ThermoFisher Scientific) using the E-Gel Ultra Low Range DNA Ladder. Amplicons were purified using the QIAquick PCR Purification Kit (Qiagen), and concentrations were quantified with the Qubit dsDNA High Sensitivity Assay Kit (ThermoFisher Scientific). Purified amplicons were pooled at equimolar concentrations and assessed for quality using a Bioanalyzer (Agilent). The final pooled library was sequenced on an Illumina NovaSeq 6000 platform at the Iowa Institute for Human Genetics (University of Iowa).

### Aptamer NGS data analysis

Raw multiplexed sequencing data in FASTQ format were uploaded to the Galaxy platform (usegalaxy.org) for barcode demultiplexing.^34^ The resulting barcode-split data were compiled into a non-redundant aptamer database using Galaxy workflows.^34^ The internal reference sequence (Control-2F) was identified, and the Sel2N25 aptamers were filtered to retain only those with variable regions between 23 and 27 nucleotides in length. The distribution of aptamer read counts across experimental replicates was assessed, and aptamers present in at least three out of four replicates within the human VSMC dataset were selected for further analysis within Microsoft Excel. Read counts for each aptamer in each replicate were normalized to reads per million (RPM), relative to the internal reference read count detected for that replicate. Specificity scores were calculated as the fold difference in normalized read counts between VSMCs and endothelial cells for both human and mouse datasets. Frequency distributions of specificity scores were generated using GraphPad Prism. Statistical significance between VSMC and endothelial cell read counts was determined using multiple unpaired t-tests in GraphPad Prism for both species’ datasets. Clustering was accomplished using AptamerRunner and visualized using Cytoscape.

### SELEX Selection Round NGS

Aptamer RNA from each SELEX selection round was prepared for NGS using the previously described Illumina reverse primer and barcoded forward primer. Sequencing data were processed and compiled into a non-redundant aptamer database for each round. To enable comparative analysis, the selection round database was joined with the ASSET NGS non-redundant database using the “Join two Datasets” function in Galaxy. Read counts for each aptamer within each selection round were normalized to reads per 10 million (RP10M), based on the total number of reads acquired for that round. Aptamers were named based on their rank order within the ninth selection round. The Log2 fold enrichment between selection rounds was calculated using these normalized read counts to determine round-to-round aptamer enrichment.

### Single aptamers

Single aptamer ssDNA template oligonucleotides were synthesized by Integrated DNA Technologies (IDT), with the first two nucleotides at the 5′ end modified with OMe as described for the Sel2N25 aptamer library. Template oligos were converted to double-stranded DNA (dsDNA) using previously published methods. The sequences of aptamer RNAs and their corresponding template oligos are listed in **Supplemental Table 1**. 13-2F, 41-2F and Control-2F aptamers were transcribed *in vitro* using 2′-fluoro-pyrimidine-modified nucleotides, following established protocols. Sel2N25 OMe VSMC aptamers (V1, V2, V4, V28, V41) and false-positive negative controls (V7, V8) incorporating 2′-OMe-pyrimidine modifications were transcribed using the same methods applied to the 2′-OMe-pyrimidine-modified RNA aptamer library.

### Data analysis and statistics

Data processing, analysis, and visualization was performed using Galaxy (usegalaxy.org), Microsoft Excel, GraphPad Prism v10.4.2, AptamerRunner, and Cytoscape v3.7.2. All statistical tests were conducted in GraphPad Prism. Associated data files and figure metadata are provided in **Supplemental Data**.

## FUNDING

This work was supported by the National Institutes of Health (R01HL139581 and R01157956 to WHT; K22CA263783 to KWT), the Department of Defense (DOD CDMRP-PRCRP CA220729 to KWT), the American Cancer Society (HCCC, IRG-18-165-43), the Bunge and Born Fund (FBB-20170609 to DRC) and Fulbright-Argentinian Ministry of Education (ME-FLB-2022-2023 to DRC).

## CONFLICT OF INTEREST

None

## Notes

### Competing Interest Statement

The authors have declared no competing interest.

