## Supplemental Figures and Tables for "ASSET: A Framework for Decoding Aptamer Specificity of an Enriched Library by Next-Generation Sequencing of Experimental Samples"

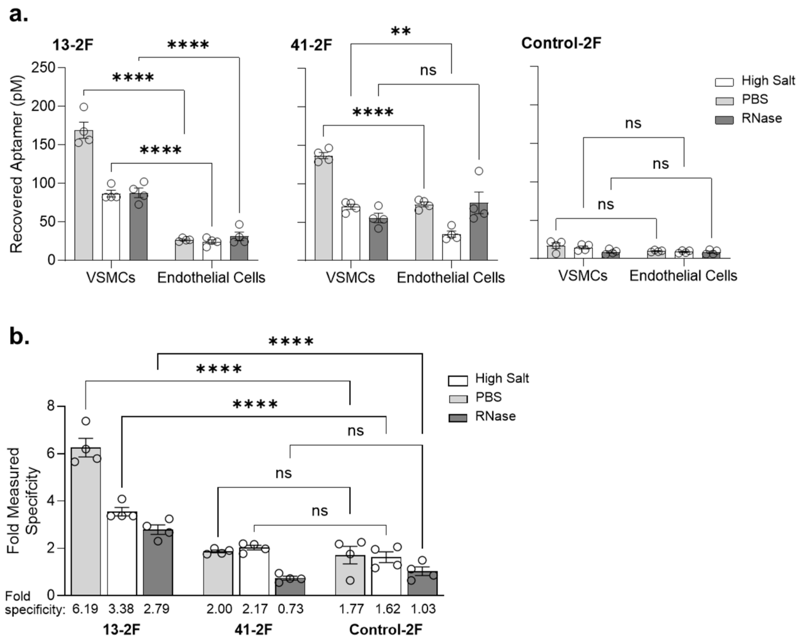

**Supplemental Figure 1: 2'-fluoro (2F) pyrimidine-modified VSMC specificity for human VSMCs.** (**a**) Quantitative PCR of recovered aptamer from human VSMCs or endothelial cells treated with 13-2F, 41-2F, or Control 2F. After incubation of aptamer, cells were treated with either PBS, high salt (0.5M NaCl), or RNase cocktail. (**b**) Fold specificity of the 2F aptamers for human VSMCs over endothelial cells. N= 4, 2-way ANOVA with multiple comparisons, ** $p < 0.01$, **** $p<0.0001$, ns = not significant.

a.

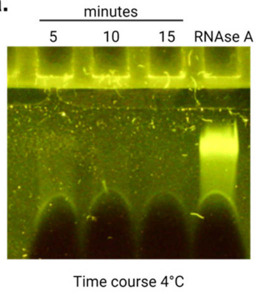

b.

| Round | Apatmer (nM) | Negative Cell | Negative Time (min.) | Positive Cell | Positive Time (min.) | RNase Cocktail |
|---|---|---|---|---|---|---|
| 1a-c | 300 | hEC x1 | 15 | hVSMC | 30 | 5 min. 4°C |
| 2a-c | 300 | hEC x1 | 30 | hVSMC | 15 | 5 min. 4°C |
| 3a-c | 300 | hEC x1 | 30 | hVSMC | 15 | 5 min. 4°C |
| 4 | 300 | mEC x1 | 30 | mVSMC | 15 | 5 min. 4°C |
| 5 | 150 | mEC x1 | 30 | mVSMC | 15 | 5 min. 4°C |
| 6 | 150 | hEC x2 | 60 | hVSMC | 15 | 5 min. 4°C |
| 7 | 75 | hEC x2 | 60 | hVSMC | 15 | 5 min. 4°C |
| 8 | 75 | mEC x2 | 60 | mVSMC | 15 | 5 min. 4°C |
| 9 | 25 | mEC x2 | 60 | mVSMC | 15 | 5 min. 4°C |

Replicates (rounds 1a-c – 3a-c)

RNase Cell-Internalization SELEX (rounds 1a-c – 9)

Cross-Species Toggle SELEX (Positive Cell)

hVSMC = human VSMCs
hEC = human endothelial cells
mVSMC = mouse VSMCs
mEC = mouse endothelial cells

**Supplemental Figure 2: SELEX conditions.** (**a**) 2'-o-methyl (OMe) pyrimidine-modified aptamer library incubated at 4°C with the RNase Cocktail for 5 – 15 minutes or RNAse A for 15 minutes. (**b**) Conditions applied during RNAse Cell-Internalization and Cross-Species Toggle SELEX to enrich for VSMC aptamers.

**a.**

| Replicate | Unique | Total | Reference |
|---|---|---|---|
| hVSMC1 | 522 | 44,477 | 4,279,068 |
| hVSMC2 | 429 | 22,898 | 4,709,918 |
| hVSMC3 | 579 | 62,171 | 4,564,794 |
| hVSMC4 | 597 | 63,585 | 4,070,889 |
| hEC1 | 300 | 3,505 | 5,102,836 |
| hEC2 | 333 | 4,580 | 5,338,918 |
| hEC3 | 316 | 4,413 | 6,215,328 |
| hEC4 | 316 | 4,413 | 6,215,328 |
| mVSMC1 | 323 | 9,744 | 4,626,024 |
| mVSMC3 | 360 | 8,508 | 4,163,441 |
| mVSMC4 | 423 | 18,643 | 4,508,147 |
| mEC1 | 291 | 2,100 | 4,047,767 |
| mEC2 | 276 | 1,705 | 4,975,093 |
| mEC3 | 262 | 2,041 | 4,675,002 |
| mEC4 | 286 | 2,288 | 5,048,981 |
| Total | 1,421 | 255,071 | 72,541,534 |

**b.**

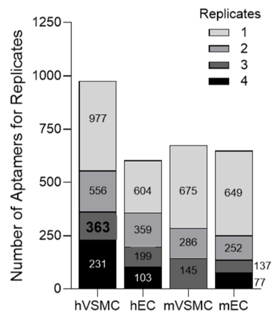

**c.**

| | Human | Mouse |
|---|---|---|
| Number of values | 115 | 115 |
| Minimum | 4.001 | 0.8522 |
| 25% Percentile | 5.793 | 1.628 |
| Median | 9.300 | 2.509 |
| 75% Percentile | 17.37 | 4.504 |
| Maximum | 40.18 | 54.78 |
| Range | 36.18 | 53.93 |
| 95% CI of median | | |
| Actual confidence level | 96.02% | 96.02% |
| Lower confidence limit | 7.570 | 2.124 |
| Upper confidence limit | 11.51 | 3.332 |
| Mean | 12.41 | 4.024 |
| Std. Deviation | 8.513 | 5.664 |
| Std. Error of Mean | 0.7939 | 0.5281 |
| Lower 95% CI of mean | 10.84 | 2.978 |
| Upper 95% CI of mean | 13.98 | 5.070 |
| Geometric mean | 10.11 | 2.842 |
| Geometric SD factor | 1.877 | 2.089 |
| Lower 95% CI of geo. mean | 9.002 | 2.480 |
| Upper 95% CI of geo. mean | 11.36 | 3.256 |

**d.**

| | Human | Mouse |
|---|---|---|
| Number of values | 22 | 22 |
| Minimum | 7.890 | 4.001 |
| 25% Percentile | 14.31 | 4.881 |
| Median | 20.13 | 6.253 |
| 75% Percentile | 24.66 | 11.07 |
| Maximum | 35.17 | 54.78 |
| Range | 27.28 | 50.78 |
| 95% CI of median | | |
| Actual confidence level | 98.31% | 98.31% |
| Lower confidence limit | 14.34 | 4.886 |
| Upper confidence limit | 24.50 | 10.73 |
| Mean | 19.85 | 9.663 |
| Std. Deviation | 7.019 | 10.69 |
| Std. Error of Mean | 1.497 | 2.279 |
| Lower 95% CI of mean | 16.74 | 4.925 |
| Upper 95% CI of mean | 22.96 | 14.40 |
| Geometric mean | 18.58 | 7.514 |
| Geometric SD factor | 1.469 | 1.840 |
| Lower 95% CI of geo. mean | 15.67 | 5.734 |
| Upper 95% CI of geo. mean | 22.04 | 9.846 |

**Supplemental Figure 3: Summary of ASSET NGS data.** (**a**) Unique and total read counts for library aptamers and internal reference across experimental samples and replicates. (**b**) Distribution of aptamers across replicates for each experimental sample. Summary statistics of the (**c**) 115 aptamers with greater than four-fold significant specificity for human VSMCs and the (**d**) 22 aptamers with greater than four-fold significant specificity for mouse VSMCs.

**a.**

| Round | Total | Unique |
|---|---|---|
| 0A | 12,294,501 | 11,155,253 |
| 0B | 8,342,721 | 7,642,670 |
| 0C | 11,538,307 | 10,547,174 |
| 1A | 9,630,376 | 4,792,545 |
| 1B | 11,433,053 | 9,830,143 |
| 1C | 12,276,269 | 10,988,323 |
| 2A | 10,627,494 | 3,789,598 |
| 2B | 10,322,099 | 6,885,986 |
| 2C | 11,824,064 | 9,315,191 |
| 3A | 5,669,329 | 2,483,234 |
| 3B | 6,157,375 | 3,742,259 |
| 3C | 423,823 | 343,059 |
| 4 | 8,405,315 | 1,818,516 |
| 5 | 8,944,566 | 565,120 |
| 6 | 7,176,828 | 90,114 |
| 7 | 6,905,527 | 84,325 |
| 8 | 8,176,251 | 85,537 |
| 9 | 7,212,123 | 92,600 |
| Total | 157,360,021 | 84,251,647 |

**b.**

Pearson's correlation

| Var 1 | Var 2 | corr | p-value |
|---|---|---|---|
| Human ASSET | Mouse ASSET | 0.72 | <0.0001 |
| Human ASSET | Round 9 RP10M | 0.18 | 0.0007 |
| Mouse ASSET | Round 9 RP10M | 0.28 | <0.0001 |
| Human ASSET | Enrichment Rnd 3 to 9 | 0.60 | <0.0001 |
| Mouse ASSET | Enrichment Rnd 3 to 9 | 0.40 | <0.0001 |

Spearman's correlation

| Var 1 | Var 2 | corr | p-value |
|---|---|---|---|
| Human ASSET | Mouse ASSET | 0.63 | <0.0001 |
| Human ASSET | Round 9 RP10M | 0.61 | <0.0001 |
| Mouse ASSET | Round 9 RP10M | 0.44 | <0.0001 |
| Human ASSET | Enrichment Rnd 3 to 9 | 0.70 | <0.0001 |
| Mouse ASSET | Enrichment Rnd 3 to 9 | 0.47 | <0.0001 |

Kendall's correlation

| Var 1 | Var 2 | corr | p-value |
|---|---|---|---|
| Human ASSET | Mouse ASSET | 0.47 | <0.0001 |
| Human ASSET | Round 9 RP10M | 0.45 | <0.0001 |
| Mouse ASSET | Round 9 RP10M | 0.31 | <0.0001 |
| Human ASSET | Enrichment Rnd 3 to 9 | 0.53 | <0.0001 |
| Mouse ASSET | Enrichment Rnd 3 to 9 | 0.34 | <0.0001 |

**c.**

| Aptamer | Abundance (Rnd 9 RP10M) | Enrichment (Log2 Rnds 3:9) | Human Specificity Score | Mouse Specificity Score |
|---|---|---|---|---|
| V1 | 3,764,803 | 3.09 | 21.29 | 12.10 |
| V2 | 2,070,113 | 3.07 | 20.22 | 15.67 |
| V3 | 1,570,963 | 2.32 | 9.49 | 2.56 |
| V4 | 643,992 | 4.42 | 21.06 | 54.78 |
| V5 | 233,623 | 3.39 | 8.72 | 2.41 |
| V6 | 150,200 | 3.69 | 6.30 | 1.57 |
| V7 | 74,806 | 1.90 | 1.41 | 1.33 |
| V8 | 47,958 | 3.87 | 3.31 | 2.78 |
| V9 | 42,417 | 1.65 | 1.43 | 1.12 |
| V10 | 41,257 | 1.70 | 1.57 | 1.23 |

**d.**

| Aptamer | Enrichment (Log2 Rnds 3:9) | Abundance (Rnd 9 RP10M) | Human Specificity Score | Mouse Specificity Score |
|---|---|---|---|---|
| V4 | 4.42 | 643,992 | 21.06 | 54.78 |
| V8 | 3.87 | 47,958 | 3.31 | 2.78 |
| V16 | 3.71 | 26,978 | 40.18 | 2.30 |
| V6 | 3.69 | 150,200 | 6.30 | 1.57 |
| V15 | 3.65 | 28,975 | 32.21 | 1.63 |
| V19 | 3.65 | 20,044 | 7.57 | 3.05 |
| V13 | 3.56 | 34,522 | 4.00 | 1.12 |
| V24 | 3.49 | 15,814 | 11.68 | 12.45 |
| V28 | 3.47 | 13,217 | 21.15 | 16.65 |
| V12 | 3.47 | 35,490 | 17.37 | 2.51 |

**Supplemental Figure 4: Summary of selection round NGS data.** (**a**) Unique and total read counts acquired from the starting aptamer libraries and selection rounds. (**b**) Correlation analysis (Pearson's, Spearman's, Kendall's) between log2 human and mouse ASSET specificity scores, aptamer abundance (Round 9 RP10M), and aptamer enrichment (Enrichment Rnd 3:9). The top ten most (**c**) abundant, and (**d**) enriched between round three to round nine in round nine.

**a.**

| | Human | Mouse |
|---|---|---|
| Number of values | 363 | 363 |
| Minimum | 0.5450 | 0.4660 |
| 25% Percentile | 1.623 | 1.051 |
| Median | 2.659 | 1.387 |
| 75% Percentile | 5.739 | 2.057 |
| Maximum | 40.18 | 54.78 |
| Range | 39.64 | 54.31 |
| 95% CI of median | | |
| Actual confidence level | 95.40% | 95.40% |
| Lower confidence limit | 2.299 | 1.122 |
| Upper confidence limit | 3.017 | 1.387 |
| Mean | 5.493 | 2.155 |
| Std. Deviation | 6.843 | 3.458 |
| Std. Error of Mean | 0.3592 | 0.1815 |
| Lower 95% CI of mean | 4.787 | 1.798 |
| Upper 95% CI of mean | 6.200 | 2.512 |
| Geometric mean | 3.330 | 1.582 |
| Geometric SD factor | 2.542 | 1.895 |
| Lower 95% CI of geo. mean | 3.025 | 1.481 |
| Upper 95% CI of geo. mean | 3.667 | 1.690 |

**b.**

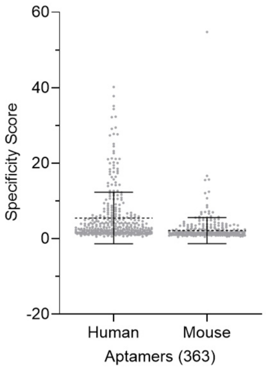

**Supplemental Figure 5: Determine threshold value for human and mouse VSMC specificity scores.** (**a**) Summary statistics and (**b**) plot (Mean ±1SD) of the 363 aptamers detected by ASSET

**a.**

| Aptamer | Human Measured Specificity | Mouse Measured Specificity | Human Specificity Score | Mouse Specificity Score | Abundance (Rnd 9 RP10M) | Enrichment (Log2 Rnds 3:9) |
|---|---|---|---|---|---|---|
| V4 | 13.20 | 71.68 | 21.06 | 54.78 | 643,992 | 4.42 |
| V28 | 44.13 | 44.72 | 21.15 | 16.65 | 13,217 | 3.47 |
| V41 | 15.14 | 27.13 | 25.13 | 15.57 | 8,172 | 3.26 |
| V2 | 24.28 | 21.65 | 20.22 | 15.67 | 2,070,113 | 3.07 |
| V1 | 19.79 | 25.05 | 21.29 | 12.10 | 3,764,803 | 3.09 |
| V7 | 1.07 | 3.77 | 1.41 | 1.33 | 74,806 | 1.90 |
| V8 | 0.71 | 2.55 | 3.31 | 2.78 | 47,958 | 3.87 |

**b.**

Pearson's correlation

| Var 1 | Var 2 | corr | p-value |
|---|---|---|---|
| Measured specificity | ASSET specificity score | 0.84 | 0.001 |
| Measured specificity | Round 9 RP10M | 0.06 | ns |
| Measured specificity | Enrichment Rnd 3 to 9 | 0.43 | ns |

Spearman's correlation

| Var 1 | Var 2 | corr | p-value |
|---|---|---|---|
| Measured specificity | ASSET specificity score | 0.57 | 0.035 |
| Measured specificity | Round 9 RP10M | -0.01 | ns |
| Measured specificity | Enrichment Rnd 3 to 9 | 0.19 | ns |

Kendall's correlation

| Var 1 | Var 2 | corr | p-value |
|---|---|---|---|
| Measured specificity | ASSET specificity score | 0.36 | ns |
| Measured specificity | Round 9 RP10M | 0.07 | ns |
| Measured specificity | Enrichment Rnd 3 to 9 | 0.21 | ns |

**Supplemental Figure 6: Candidates and controls.** (**a**) Summary of human and mouse measured specificity, human and mouse ASSET specificity scores, and selection round NGS data for candidates and controls. (**b**) Correlation analysis between measured specificity, ASSET specificity scores, aptamer abundance (Round 9 RP10M), and aptamer enrichment (Enrichment Rnd 3 to 9). ns = not significant.

### SUPPLEMENTAL TABLES

#### Supplemental Table 1: Aptamer sequences and template oligos.

**2’-fluoro-pyrimidine modified VSMC aptamers**

13-2F
Aptamer RNA : 5'-GGGAGGACGAUGCGGUGAGUCGUUCCCUUCGUCCCCAGACGACUCGCCCGA-3'
Template Oligo: 5’-mUmCGGGCGAGTCGTCTGGGGACGAAGGGAACGACTCACCGCATCGTCCTCCC-3’

41-2F
Aptamer RNA: 5'-GGGAGGACGAUGCGGUGACUCGUCUGUUUCGUCCCCAGACGACUCGCCCGA-3'
Template Oligo: 5’-mUmCGGGCGAGTCGTCTGGGGACGAAACAGACGAGTCACCGCATCGTCCTCCC-3’

Control-2F (ASSET internal reference)
Aptamer RNA: 5’-GGGAGGACGAUGCGGAUUACGAGCUUGUUGCCUCGCAGACGACUCGCCCGA-3’
Template Oligo: 5’-mUmCGGGCGAGTCGTCTGCGAGGCAACAAGCTCGTAATCCGCATCGTCCTCCC-3’

**2’-OMe-pyrimindine modified VSMC aptamers and ASSET false positive negative controls**

*Redacted until manuscript accepted for publication*

mU mC = ssDNA template oligos synthesized with the first two nucleotides OMe-modified to minimize single nucleotide addition by the T7 RNAP during *in vitro* RNA transcription.
